# Benchmarking large language models for genomic knowledge with GeneTuring

**DOI:** 10.1101/2023.03.11.532238

**Authors:** Xinyi Shang, Xu Liao, Zhicheng Ji, Wenpin Hou

## Abstract

Large language models (LLMs) show promise in biomedical research, but their effectiveness for genomic inquiry remains unclear. We developed GeneTuring, a benchmark consisting of 16 genomics tasks with 1,600 curated questions, and manually evaluated 48,000 answers from ten LLM configurations, including GPT-4o (via API, ChatGPT with web access, and a custom GPT setup), GPT-3.5, Claude 3.5, Gemini Advanced, GeneGPT (both slim and full), BioGPT, and BioMedLM. A custom GPT-4o configuration integrated with NCBI APIs, developed in this study as SeqSnap, achieved the best overall performance. GPT-4o with web access and GeneGPT demonstrated complementary strengths. Our findings highlight both the promise and current limitations of LLMs in genomics, and emphasize the value of combining LLMs with domain-specific tools for robust genomic intelligence. GeneTuring offers a key resource for benchmarking and improving LLMs in biomedical research.

**Biographical Note:** Dr. Wenpin Hou is an Assistant Professor (tenure-track) in the Department of Biostatistics at Columbia University and member of its Data Science Institute, developing AI and statistical methods for decoding gene regulatory programs from single-cell and spatial multiomics data.

## Introductions

Large language models (LLMs), such as GPT-3.5^1^, GPT-4^2^, Gemini^3^, and Claude^4^, are advanced models trained on massive datasets, capable of producing text that closely resembles human speech. LLMs excel in various tasks, such as answering questions^2^, generating programming code^5^, and analyzing images^6^. Recent studies have also highlighted their strong capabilities in genomic research. For instance, in single-cell RNA-seq data, GPT-4 can produce cell type annotations that are highly consistent with those provided by human experts, using only marker gene information as input^7^. Additionally, gene embeddings generated by GPT-3.5 can be utilized to create single-cell embeddings for various downstream analyses^8^.

These studies suggest that LLMs possess knowledge in the field of genomics and have the potential to serve as a knowledge base for genomic research. Such an LLM-based genomic knowledge base could significantly benefit genomic research by reducing the time required to locate and retrieve reliable information, a process that is often time-consuming for interdisciplinary researchers with limited genomic expertise. Moreover, the advanced reasoning and analytical capabilities of modern LLMs enable efficient synthesis of information from diverse sources. However, whether LLMs can reliably serve as genomic knowledge bases has not been systematically studied and remains poorly understood.

Benchmark datasets are essential for comparing and assessing the ability of LLMs to perform specific tasks. For example, MMLU (Massive Multitask Language Understanding)^9^ is a widely used benchmark dataset for evaluating LLMs’ interdisciplinary knowledge, while HumanEval^10^ assesses their ability to generate programming code. These benchmark datasets provide a standardized framework for comparing performance across different LLMs and tracking model evolution over time. They have been pivotal in identifying the weaknesses of existing models and guiding future development toward more advanced LLMs. However, the benchmarks for the genomics are still lacking. A recent study^11^ assessed LLMs in phenotype-driven gene prioritization, revealing limitations in their diagnostic accuracy for rare genetic diseases. BioCoder^12^ was introduced as a benchmark for evaluating LLMs in bioinformatics code generation, underscoring the benefits of domain-specific fine-tuning. Furthermore, expansive applications of LLMs in bioinformatics were reviewed^13^, emphasizing their transformative role in tasks ranging from sequence analysis to functional annotation. These studies collectively highlight the necessity for specialized benchmarks to assess and enhance LLM performance in genomics and bioinformatics.

To this end, we developed GeneTuring, a comprehensive *knowledge-based* question-and-answer (Q&A) database, to benchmark the performance of LLMs in genomics. GeneTuring comprises 16 modules that cover a wide range of genomic research topics, including gene and single nucleotide polymorphism (SNP) locations, genomic functions, sequence alignment, and other essential areas of genomic knowledge. We evaluated the performance of ten LLM settings on GeneTuring, including BioGPT^14^, BioMedLM^15^, GPT-3.5^1^, GPT-4o^16^, GPT-4o (web), Gemini Advanced^3^, Claude 3.5^4^, GeneGPT (both slim and full)^17^, and SeqSnap (a custom GPT-4o configuration integrated with NCBI APIs, developed in this study). Among these, GPT-4o, Gemini Advanced, and Claude 3.5 are contemporary LLMs settings widely recognized for their strong performance across various tasks.

Our analysis revealed significant variation in the accuracy of genomic knowledge-based question answering across the LLMs. Moreover, we observed that performance could be further enhanced when LLMs have access to web browsing capabilities or have been integrated with domain-specific database. However, even the best-performing LLM completely failed in certain tasks and were unable to answer all questions correctly, despite the likelihood that the genomic knowledge was included in their training corpora. These findings suggest that current LLMs fall short of serving as reliable knowledge bases for genomic research due to issues such as AI hallucination^18,19^. More advanced LLMs will be required to address these limitations in the future.

## Results

GeneTuring consists of 16 modules containing a total of 1,600 question-answer pairs, which are grouped into four categories: nomenclature, genomic location, functional analysis, and sequence alignment. These modules represent tasks commonly encountered in genomic research. Figure 1 illustrates the categories and names of the 16 modules, along with example question-answer pairs, example responses, and the corresponding scores of these responses. To ensure compatibility with models based on GPT-2, the same questions were reformulated as sentence completion tasks (Methods).

**Figure 1.**
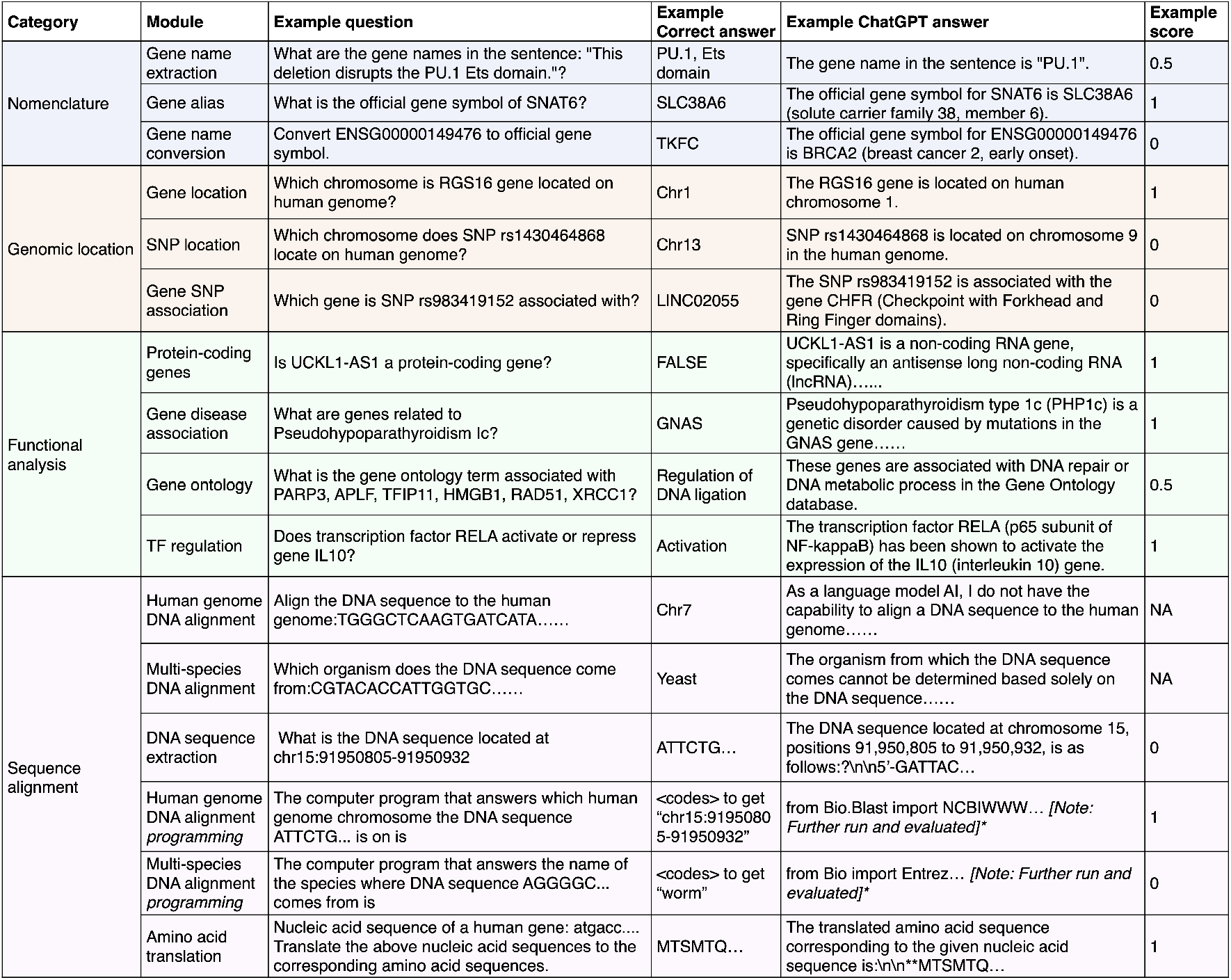
GeneTuring modules. A table summarizing all 16 GeneTuring modules, organized into four categories. Each module entry includes its name, an example question-answer pair, an example response, and the corresponding evaluation score.

We used GeneTuring to evaluate ten LLM settings: BioGPT^14^, BioMedLM^15^, Claude 3.5^4^, GeneGPT^17^ (both slim and full), Gemini Advanced^3^, GPT-3.5^1^, GPT-4o^16^, GPT-4o (web) (i.e., GPT-4o with web access), and SeqSnap (a custom GPT-s application developed in this study). GPT-3.5, GPT-4o, and GPT-4o (web) are commercial models developed by OpenAI. SeqSnap is a custom application developed through GPT Builder platform and connected with NCBI APIs. Gemini Advanced and Claude 3.5 are also commercial models, developed by Google and Anthropic, respectively. BioGPT and BioMedLM are based on the GPT-2 architecture but are specifically trained on biomedical literature. GeneGPT is based on GPT3.5-turbo and implemented in both slim and full versions. We accessed all models via API without web access, except for GPT-4o (web), which was evaluated through the web browser version to assess whether web access influences performance. GeneGPT and SeqSnap were included to evaluate whether access to external databases can improve performance on sequence analysis tasks.

For BioGPT and BioMedLM, each question was presented once, and the top three answers returned by the model were recorded. For all other models, each question was presented three times and the three responses were recorded. For each of the 48,000 responses, we manually evaluated whether each model correctly understood the questions (comprehending capacity), provided seemingly correct but actually false answers (AI hallucination), acknowledged its inability to answer the question (incapacity awareness), or correctly returned answers (accuracy). If the question was understood, a numeric score between 0 and 1 was assigned based on a predefined scoring mechanism, unless the LLM acknowledged its incapacity (Methods). All Q&A pairs used in the evaluation, responses from the LLMs, and the corresponding scores are provided on our project website https://github.com/Winnie09/GeneTuring.

We first assessed whether the LLMs could understand the questions and generate relevant responses (Figure 2). GPT-4o, SeqSnap, Claude 3.5, and Gemini Advanced consistently provided relevant answers to all questions. GPT-4o (web), GPT-3.5, and GeneGPT (both slim and full) correctly understood nearly all questions. In contrast, GPT-2-based models (BioGPT and BioMedLM) struggled to interpret many tasks, probably due to their limited model capacity and reliance on a sentence-completion mechanism. As understanding the question is a prerequisite for generating correct answers, these results emphasize the importance of using more advanced model architectures with stronger natural language understanding capabilities.

**Figure 2.**
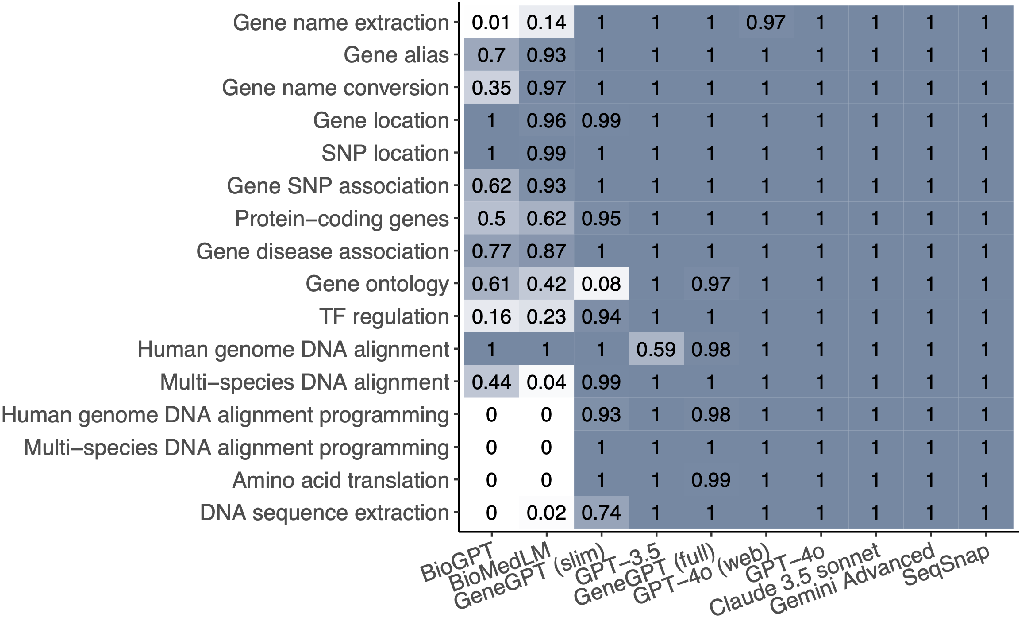
Comprehending capacity. The proportion of questions that were correctly understood by each method in each module. Methods (columns, left to right) are ordered by increasing average proportion.

We then evaluated the extent of AI hallucination which was defined as the generation of confident but inaccurate answers despite correctly understanding the question (Figure 3). This was measured as the proportion of questions that received a score of zero for their responses. Hallucination was observed across models and tasks, including in advanced LLMs such as GPT-4o and GPT-3.5. For example, GPT-4o produced incorrect answers to almost all questions related to SNP locations. Additionally, all models exhibited high levels of hallucination in the gene name conversion task except GeneGPT (slim) and GPT-4o (web).

**Figure 3.**
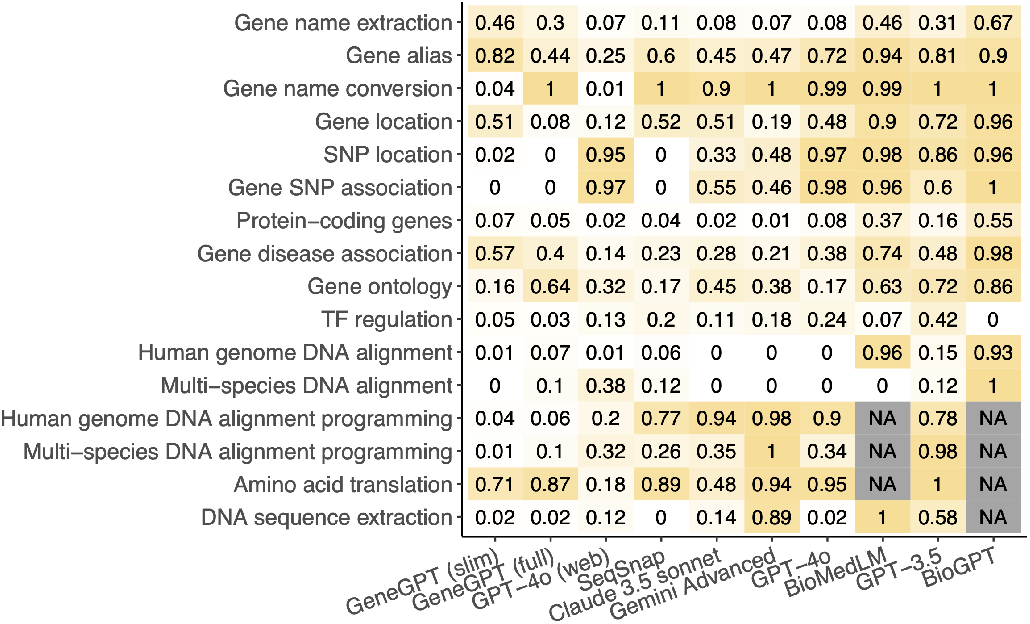
AI hallucination degree. Among the questions that were correctly understood, the proportion of answers with zero scores. Methods (columns, left to right) are ordered by increasing average proportion.

Results indicate that access to online content is an effective strategy for improving accuracy, as demonstrated by the performance difference between GPT-4o and GPT-4o (web) on gene name conversion tasks. While GPT-4o without web access produced errors in 99% of cases (Figure 3), GPT-4o with browsing capabilities achieved 99% accuracy (Figures 3, 5). A similar improvement was observed in the gene alias and gene location tasks, where error rates dropped substantially when GPT-4o had access to the web. GPT-4o (web) was the only model to achieve high accuracy in gene name conversion (99%) and outperformed all others on gene alias and gene location queries, benefiting from access to online resources. However, online access is not universally beneficial across all tasks. For example, GPT-4o’s performance on two SNP-related tasks remained largely unchanged regardless of web access. A likely explanation is that gene names and IDs are widely available in online databases and literature, while many SNPs represent rare variants that are sparsely documented or not publicly available online. Therefore, LLMs benefit from online access only when relevant information exists within accessible web resources.

We further investigated why AI hallucination is less severe in certain cases. We found that LLMs are sometimes able to recognize and report their inability to answer a question without providing a definite response, a phenomenon that we refer to as “incapacity awareness”. For example, in one of the multi-species DNA alignment tasks, Gemini Advanced responded, “Unfortunately, determining the exact organism from this short DNA sequence alone is not possible.” We evaluated incapacity awareness using the proportion of cases where LLMs reported their incapacity to answer, given that the question is properly understood (Figure 4). This proportion is particularly high in the two tasks related to DNA sequence alignment, explaining the relatively low occurrence of AI hallucinations in these tasks. In contrast, as expected, the proportion of incapacity awareness is low in tasks such as gene name conversion, where AI hallucination is more prevalent.

**Figure 4.**
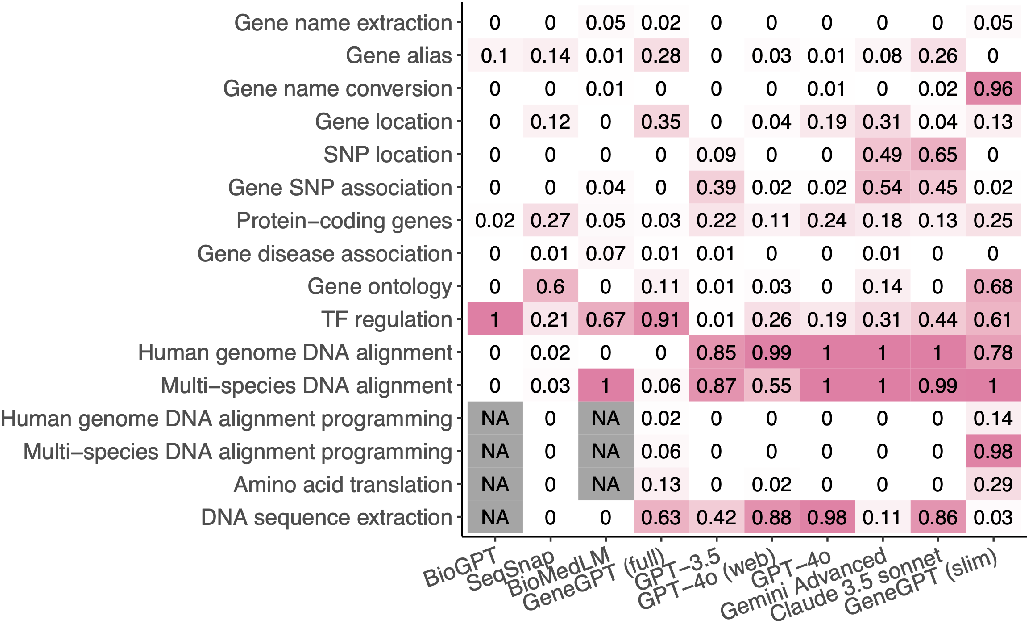
Incapacity awareness. The proportion of questions where the model acknowledged its incapacity in answering the question, calculated only for questions that were correctly understood. Methods (columns, left to right) are ordered by increasing average proportion.

**Figure 5.**
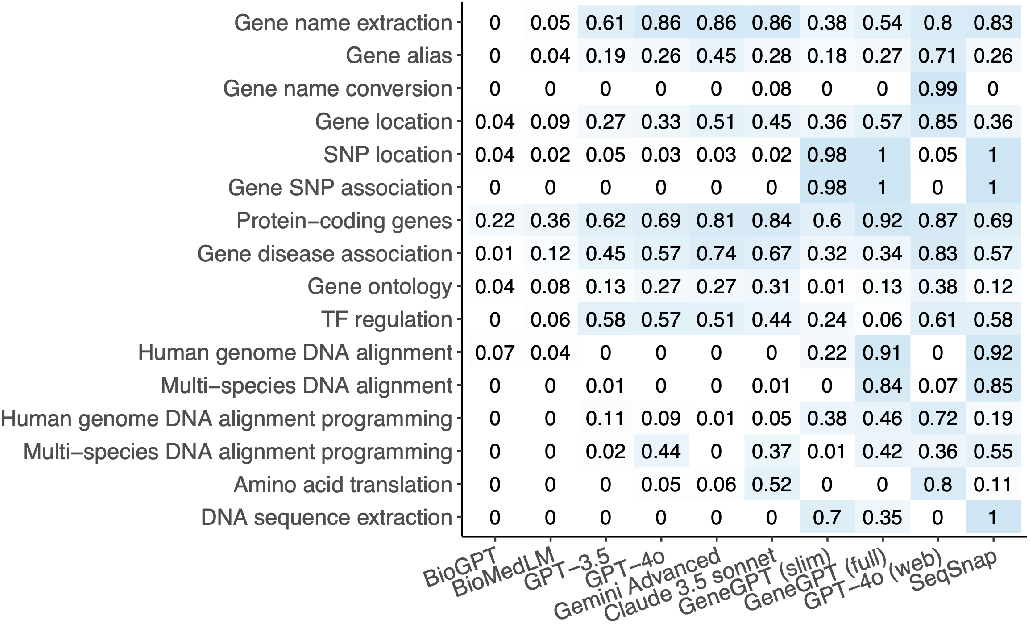
Accuracy. Among all questions, this metric represents the proportion receiving correct or partially correct answers (i.e., scores of 1 or 0.5). Methods (columns, left to right) are ordered by increasing average proportion.

We argue that incapacity awareness is crucial for addressing AI hallucination, particularly in scientific research areas such as genomics. Each version of an LLM is trained on a finite dataset, and inevitably, there will be questions that fall outside the scope of its training data, making it unlikely for the model to provide correct answers. Admitting these limitations alerts users to seek alternative approaches. In contrast, when LLMs generate random or false answers, users are often unaware of the inaccuracies, which can lead to further issues, such as wasted time and resources on validation experiments for incorrect targets.

The accuracy summary highlights that most LLMs are unable to perform sequence analysis tasks (Figure 5). The primary reason is their lack of built-in access to the complete human reference genome and related genome databases. Commercial models such as ChatGPT, Gemini Advanced, and Claude 3.5 Sonnet do not disclose their training data, but based on their outputs, it is evident that reference genome data (e.g., GRCh38) is not included in their environments. As a result, these models cannot accurately align DNA sequences to the genome or retrieve sequences from specified regions.

To address this limitation, we explored two alternative strategies. First, LLMs can be trained to interface with external genomic databases, such as the National Center for Biotechnology Information (NCBI), instead of relying solely on their internal training data, as demonstrated by GeneGPT (Figure 5). While other models showed near-zero accuracy, GeneGPT (full) achieved excellent performance in human genome DNA alignment (91%) and multi-species DNA alignment (84%). The variation between GeneGPT (full) and GeneGPT (slim) is likely due to the instability of GPT-3.5-turbo and its sentence-completion-based architecture. Its amino acid translation accuracy remained at 0%, which was expected as GeneGPT’s designed prompts were not specifically optimized for this task. To demonstrate the effectiveness of API-driven execution for sequence alignment and the benefits of leveraging a more advanced language model, we developed a custom GPT-4o-based application, SeqSnap. SeqSnap integrates the NCBI BLAST and Entrez Programming Utilities (E-utilities) APIs, enabling GPT-4o to classify user queries, submit BLAST and E-utilities requests, monitor their progress, and retrieve alignment results. We implemented both API-integrated and conversational interfaces for SeqSnap, which showed comparable performance across sequence analysis, gene nomenclature, and genomic location tasks (Figure S1). The results presented here primarily reflect the API-integrated implementation. Second, we evaluated LLMs’ ability to generate executable code for sequence alignment. Two programmatic modules were designed: one for human genome DNA alignment and another for multi-species alignment. The generated code was executed and assessed for accuracy.

SeqSnap demonstrated outstanding performance, achieving 92% accuracy in human genome alignment and 85% in multi-species alignment (Figure 5). Notably, only SeqSnap and GeneGPT could perform DNA alignment; all other models either returned near-zero accuracy (e.g., BioMedLM, BioGPT, GPT-3.5, Claude) or reported inability to complete the task. GPT-4o (web) and SeqSnap successfully generated executable code for both DNA sequence and multi-species alignment, whereas GeneGPT (slim) could only get answers for human DNA alignment. GPT-4o (web) and Claude achieved the highest accuracy for amino acid translation. Notably, SeqSnap and GeneGPT were the only models capable of performing DNA sequence extraction, with 100% and 70% (or 35% in full model) accuracy, respectively. In fact, GeneGPT (slim)’s lower accuracy was due to the lower comprehending capacity (74%, Figure 2).

Performance on the programming modules indicated that most models exhibited strong understanding of programming logic and syntax, with the exception of BioGPT and BioMedLM (Figure 2). GPT-4o and SeqSnap showed the highest accuracy in generating correct code for alignment tasks. GeneGPT (slim) and Claude 3.5 Sonnet performed well on one of the alignment tasks but failed the other. GPT-3.5, GPT-4o, and Gemini Advanced often produced incorrect or non-functional code (Figures 3, 5). Common errors included misuse of genomic libraries, incorrect handling of biological data structures, and hallucinated functions, contributing to low scores in incapacity awareness (Figure 4). Despite their strong language capabilities, these models displayed notable inconsistencies in code reliability, underscoring the need for domain-specific grounding and execution validation when applying LLMs to computational genomics tasks (Figure 5).

We reported the overall score for each LLM, calculated as the average score of all responses within each module (Figure 6). To penalize AI hallucination, responses that were entirely incorrect (original score of 0) were reassigned a score of −1. A score of 0 was assigned when the model either failed to understand the question or explicitly acknowledged its incapacity. SeqSnap achieved the highest overall performance, followed by GPT-4o (web) and GeneGPT. SeqSnap answered all questions correctly in the SNP location, gene–SNP association, and DNA sequence extraction tasks, and performed well across many other tasks. Its particularly high accuracy on sequence alignment and extraction tasks was attributed to its integration of GPT-4o with the NCBI APIs. While GPT-4o (web) demonstrated strong performance, it cannot perform most of the sequence-related tasks, highlighting the limitations of even the most advanced LLMs when not connected to a domain-specific database. GPT-4o’s suboptimal performance on tasks involving SNPs and DNA sequences was likely due to the absence of such information in its training data and the limited availability of these data online. Among models without web access, GPT-4o, Claude 3.5, and Gemini Advanced performed comparably. However, their performance was substantially lower than that of GPT-4o (web) on tasks such as gene alias resolution, gene name conversion, and gene location. In contrast, BioGPT, BioMedLM, and GPT-3.5, based on earlier LLM architectures, exhibited the poorest overall performance.

**Figure 6.**
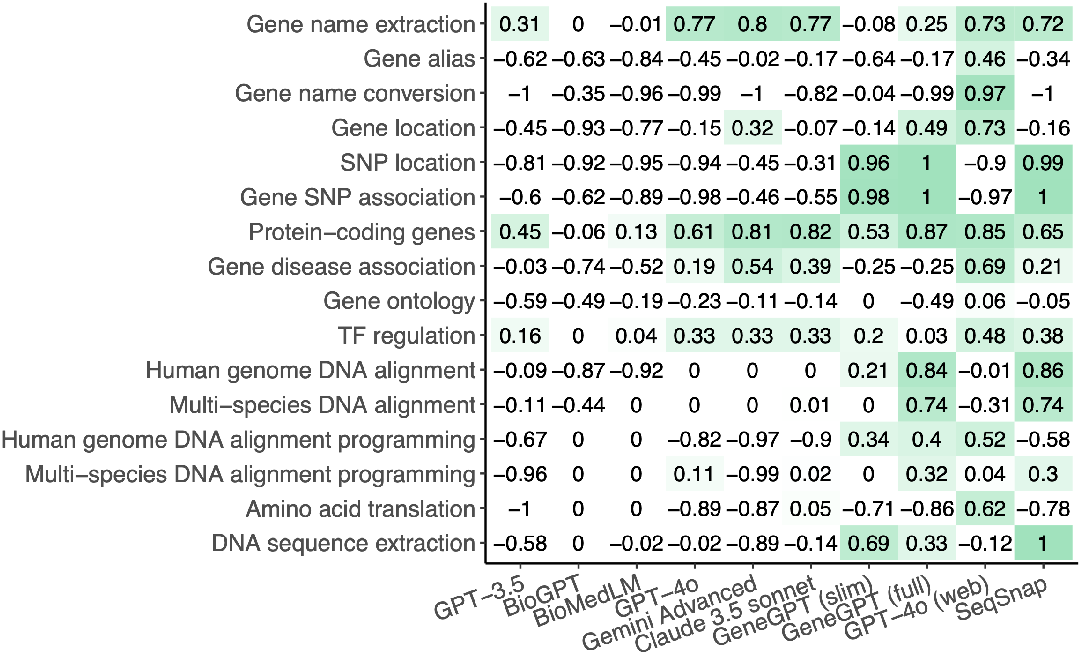
Overall score. Heatmap displaying the average score of all answers within each module. Methods (columns, left to right) are arranged in ascending order of their overall average scores.

## Conclusions

We developed GeneTuring, a first-of-its-kind Q&A benchmarking database to systematically evaluate LLMs on knowledge-based question answering in genomics. Our results reveal that while general-purpose LLMs such as GPT-4o and Claude exhibit promising capabilities, they often lack domain-specific precision and occasionally generate incorrect or unverifiable information. These limitations reflect the mismatch between the training objectives of general LLMs and the factual demands of biomedical applications. Therefore, precautions should be taken when using LLMs to answer genomics-related questions.

Improving model performance in genomics requires targeted strategies. A promising direction is tool augmentation. Integrating with domain-specific databases such as NCBI, as demonstrated in SeqSnap and GeneGPT, provides an excellent performance improvement in sequence analysis and genomic location tasks. The extension of GPT-3.5 or GPT-4 with access to NCBI Web APIs enables the model to retrieve real-time data from resources such as Entrez and BLAST during inference. This strategy significantly reduces hallucinations and improves accuracy. Such results demonstrate that coupling LLMs with structured, trusted databases can greatly improve their reliability for genomics question answering. In addition, fine-tuning on curated genomics corpora, such as gene annotations, biomedical literature, and structured databases, can help LLMs internalize more relevant knowledge. Additional gains may come from incorporating domain-specific ontologies, multi-modal inputs (e.g., sequence and expression data), or contrastive learning objectives tailored to biological reasoning. We also recognize the value of complementary benchmarks such as BioCoder^12^, which focus on LLMs’ ability to perform coding tasks in bioinformatics. As future directions, we envision integrating such programmatic evaluation frameworks into GeneTuring to provide a more holistic assessment of LLMs’ capabilities across knowledge retrieval and practical execution in genomics.

Together, these strategies, including domain-specific tool integration and fine-tuning, as well as programmatic evaluation, offer a path forward for building more accurate, interpretable, and useful language models for genomics research.

## Methods

### GeneTuring compilation and scoring criteria

GeneTuring comprises 16 modules, each containing 100 pairs of questions and answers. Additionally, GeneTuring converts each question-and-answer pair into a sentence completion task with the same meaning to accommodate models built on GPT-2 architectures. The details of how these 16 modules were compiled and scored numerically are discussed below. Notably, only relevant and definitive answers from LLMs were scored numerically. Outputs from LLMs were not scored numerically if they failed to directly answer the questions or acknowledged their limitations in doing so.

### Gene name extraction

This module evaluates the ability of LLMs to extract gene and gene product names from a given sentence. We downloaded the test set for the BioCreative II Challenge Task 1A: Gene Mention Tagging (BC2GM) from the BioCreative website^20^. The test set consists of 5,000 pairs of sentences and the corresponding names of genes and gene products mentioned in those sentences. From this set, 100 pairs were randomly selected.

To create question-answering tasks, a question was generated by appending the phrase “What are the gene and protein names in the sentence:” before the BC2GM sentence and adding a question mark at the end. For sentence completion tasks, a prompt was created by appending the phrase “The gene and protein names in the sentence” before the BC2GM sentence and adding “ is” at the end. The gene and gene product names provided by the original BC2GM task served as the gold standard answers. Each sentence may contain zero, one, or multiple gene and gene product names.

A Jaccard index was used to evaluate the performance of the answers provided by an LLM. Let 𝔸 represent the set of gene and gene product names identified by the LLM, and 𝔹 represent the gold standard set of gene and gene product names. The Jaccard index is calculated as 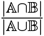, where |.| denotes the cardinality of a set. If both the gold standard and the LLM report no gene or gene product names ( 𝔸 = ∅ and 𝔹 = ∅), the Jaccard index is set to 1.

### Gene alias

This module evaluates the ability of LLMs to identify the official gene symbol for a given gene alias. The information on official gene names and their aliases for human protein-coding genes was downloaded from the NCBI website^21^. A total of 100 genes with at least one alias were randomly selected. For each gene, one alias was randomly chosen to generate the question. To create question-answering tasks, a question was generated by appending “What is the official gene symbol of” before the alias and adding a question mark after it. For sentence completion tasks, a prompt was created by appending “The official gene symbol of gene” before the alias and adding “ is” after it. The official gene symbol was used as the gold standard. Similar to the previous section, a Jaccard index was used to compare the official gene symbols provided by an LLM with the gold standard and to assign a score. The Jaccard index was employed because LLMs may provide multiple official gene symbols as answers to some questions.

### Gene name conversion

This module evaluates the ability of LLMs to convert Ensembl gene names into gene symbols. The gene annotation GTF file for the human GRCh38 genome was downloaded from the GENCODE website^22^. A total of 100 protein-coding genes were randomly selected, and the Ensembl gene name and corresponding gene symbol for each gene were recorded. To create question-answering tasks, a question was generated by appending “Convert” before the Ensembl gene name and “ to official gene symbol.” after it. For sentence completion tasks, a prompt was created by appending “The official gene symbol of” before the Ensembl gene name and “ is” after it. The gene symbol was used as the gold standard. A score of 1 was assigned if the gene symbol provided by the LLM matched the gold standard, and a score of 0 was assigned if it did not.

### Gene location

This module evaluates the ability of LLMs to determine which chromosome a gene is located on. The gene annotation GTF file for the human GRCh38 genome was downloaded from the GENCODE website^22^. Genes with multiple locations on the genome were excluded. A total of 100 genes were randomly selected, and for each gene, its gene symbol and the chromosome name it is located on were recorded. To create question-answering tasks, a question was generated by appending “Which chromosome is” before the gene symbol and “ gene located on the human genome?” after it. For sentence completion tasks, a prompt was generated by appending “ gene is located on human genome chromosome” after the gene symbol. The chromosome name was used as the gold standard. A score of 1 was assigned if the chromosome name provided by the LLM matched the gold standard, and a score of 0 was assigned if it did not.

### SNP location

This module evaluates the ability of LLMs to identify which chromosome a single nucleotide polymorphism (SNP) is located on. The SNP locations were downloaded from the NCBI website^21^. A total of 100 SNPs were randomly selected. For each SNP, its symbol and the name of the chromosome it is located on were recorded. To create question-answering tasks, a question was generated by appending “Which chromosome does SNP” before the SNP symbol and “ locate on the human genome?” after it. For sentence completion tasks, a prompt was generated by appending “SNP” before the SNP symbol and “ is located on human genome chromosome” after it. The chromosome name was used as the gold standard. A score of 1 was assigned if the chromosome name provided by the LLM matched the gold standard, and a score of 0 was assigned if it did not.

### Gene SNP association

This module evaluates the ability of LLMs to identify which gene a single nucleotide polymorphism (SNP) is associated with. SNP-gene association information was downloaded from the NCBI website^21^. SNPs associated with zero or more than one gene were excluded. A total of 100 SNPs were randomly selected, and for each SNP, its symbol and the associated gene symbol were recorded. To create question-answering tasks, a question was generated by appending “Which gene is SNP” before the SNP symbol and “ associated with?” after it. For sentence completion tasks, a prompt was generated by appending “The name of the gene associated with SNP” before the SNP symbol and “ is” after it. The gene symbol was used as the gold standard. A score of 1 was assigned if the gene symbol provided by the LLM matched the gold standard, and a score of 0 was assigned if it did not.

### Protein-coding genes

This module evaluates the ability of LLMs to determine whether a gene codes for a protein. Gene information was downloaded from the NCBI website^21^. A total of 100 human genes, including protein-coding genes, non-coding RNA (ncRNA) genes, and pseudogenes, were randomly selected. For each gene, its symbol and gene type were recorded. To create question-answering tasks, a question was generated by appending “Is” before the gene symbol and “ a protein-coding gene?” after it. For sentence completion tasks, a prompt was generated by appending “Regarding whether the gene codes for a protein,” before the gene symbol and “ is” after it. A binary value indicating whether the gene is a protein-coding gene was used as the gold standard. A score of 1 was assigned if the LLM’s response matched the gold standard in determining whether the gene is a protein-coding gene, and a score of 0 was assigned if it did not.

### Gene disease association

This module evaluates the ability of LLMs to identify genes associated with specific diseases. Gene-disease association information was downloaded from the OMIM website^23^. A total of 100 diseases were randomly selected, and all genes associated with each disease were recorded. To create question-answering tasks, a question was generated by appending “What are the genes related to” before the disease name and adding a question mark after it. For sentence completion tasks, a prompt was generated by appending “The name of the gene related to” before the disease name and “ is” after it. The set of genes associated with each disease was used as the gold standard. The answer provided by an LLM was scored based on the proportion of gold standard genes mentioned in the response.

### Gene ontology

This module evaluates the ability of LLMs to identify gene ontology (GO) terms enriched in a set of genes. GO information for biological processes was downloaded from MSigDB^24^. A total of 100 GO terms were randomly selected, and all genes associated with each GO term were recorded. To create question-answering tasks, a question was generated by appending “What is the enriched gene ontology term associated with” before the list of genes and adding a question mark after the list of genes. For sentence completion tasks, a prompt was generated by appending “The enriched gene ontology term associated with” before the list of genes and “ is” after the list of genes. The name of the GO term was used as the gold standard. A score of 1 was assigned if one of the GO terms provided by an LLM fully matched the gold standard. A score of 0.5 was assigned if one of the GO terms partially matched the gold standard. A score of 0 was assigned if none of the GO terms provided by the LLM fully or partially matched the gold standard.

### TF regulation

This module tests the ability of LLMs to determine whether a transcription factor activates or represses the expression of a gene. Information on transcription factors regulating genes in humans was obtained from the Trrust database^25^. A total of 100 pairs of transcription factors and genes with known activation or repression relationships were randomly selected. To create question-answering tasks, questions were generated by combining the phrases “Does transcription factor,” the name of the transcription factor, “activate or repress gene,” the name of the gene, and “?” into a single sentence. Similarly, to create sentence completion tasks, sentences were generated by combining: “The regulatory relationship between transcription factor,” the name of the transcription factor, “and gene,” the name of the gene, and “is” into a single prompt. A binary value indicating whether the transcription factor activates the gene served as the gold standard. A score of 1 was assigned if the LLM’s response agreed with the gold standard in identifying whether the relationship was activation or repression, while a score of 0 was assigned if they disagreed.

### Human genome DNA alignment

This module evaluates the ability of LLMs to identify the chromosome to which a DNA sequence aligns in the human genome. DNA sequence information for the human genome was obtained from the Bioconductor package BSgenome.Hsapiens.UCSC.hg38^26^. Only autosomes and sex chromosomes were retained, and regions containing “N” in the DNA sequence were excluded.

To select a genomic region, a chromosome was randomly chosen, along with a starting position within the range of the selected chromosome and a length between 100 and 150 base pairs. The corresponding DNA sequence was extracted based on the selected chromosome, starting position, and length. This process was repeated 100 times to generate 100 DNA sequences and their associated chromosome names. For question-answering tasks, questions were generated by appending “Align the DNA sequence to the human genome:” before the DNA sequence. For sentence completion tasks, questions were created by appending “The DNA sequence” before the DNA sequence and “is on the human genome chromosome” after the DNA sequence. The chromosome name was used as the gold standard. A score of 1 was assigned if the chromosome name provided by the LLM matched the gold standard, and a score of 0 was assigned if it did not.

### Multi-species DNA alignment

This module evaluates the ability of LLMs to identify the species from which a DNA sequence originates. DNA sequence information for human, mouse, rat, chicken, zebrafish, worm, and yeast genomes was obtained from the Bioconductor^26^ pack-ages, including BSgenome.Hsapiens.UCSC.hg38, BSgenome.Mmusculus.UCSC.mm10, BSgenome.Rnorvegicus.UCSC.rn5, BSgenome.Ggallus.UCSC.galGal6, BSgenome.Drerio.UCSC.danRer11, BSgenome.Celegans.UCSC.ce11, and BSgenome. Scerevisiae.UCSC.sacCer1. Only autosomes and sex chromosomes were retained, and regions containing “N” in the DNA sequence were excluded.

To select a genomic region, one of the seven species was randomly chosen, followed by the random selection of a chromosome within the chosen species, a starting position within the range of the selected chromosome, and a length between 100 and 150 base pairs. The corresponding DNA sequence was extracted based on the selected species, chromosome, starting position, and length. This process was repeated 100 times to generate 100 DNA sequences along with their respective species names. For question-answering tasks, questions were created by appending “Which organism does the DNA sequence come from:” before the DNA sequence. For sentence completion tasks, questions were created by appending “The name of the species where the DNA sequence” before the DNA sequence and “comes from is” after the DNA sequence. The species name served as the gold standard. A score of 1 was assigned if the species name provided by the LLM matched exactly with the gold standard. A score of 0.5 was assigned if the LLM provided a correct superset of the gold standard but did not match it exactly. A score of 0 was assigned if the LLM’s answer was neither a superset nor an exact match with the gold standard.

### Human genome DNA alignment programming

This module evaluates the ability of LLMs to generate code that identifies the chromosome to which a given DNA sequence aligns in the human genome. The DNA sequences and genomic regions used in this module are the same as those in the *Human genome DNA alignment* module. To generate question-answering tasks, we created prompts by prepending one of the following instructions to each DNA sequence: “Generate programming code to align the DNA sequence to the human genome:” (for GPT-4 (web)) or “Only generate Python programming code, do not provide any explanation to align the DNA sequence to the human genome:” (for all other models to avoid irrelevant text outputs in addition to codes). The correct chromosome name was used as the gold standard. A score of 1 was assigned if the chromosome name returned by the LLM-generated code matched the gold standard. A score of 0.5 was given if the code was executable and accessed relevant databases or performed alignment but failed to produce an exact match. A score of 0 was assigned if the code was not executable or produced irrelevant results.

### Multi-species DNA alignment programming

This module evaluates the ability of LLMs to generate code that identifies the species from which a given DNA sequence originates. The DNA sequences and genomic regions used here are the same as those in the *Multi-species DNA alignment* module. Prompts were generated by prepending one of the following instructions to each DNA sequence: “Generate programming code to answer which organism the DNA sequence comes from:” (for GPT-4(Web)) or “Only generate Python programming code, do not provide any explanation to answer which organism the DNA sequence comes from:” (for all other models to avoid irrelevant texts outputs in addition to codes). The species name served as the gold standard. A score of 1 was assigned if the species name returned by the LLM-generated code matched the gold standard. A score of 0.5 was given if the code was executable and accessed relevant databases or performed alignment but failed to produce an exact match. A score of 0 was assigned if the code was not executable or produced irrelevant results.

### Amino acid translation

This module evaluates the ability of LLMs to translate nucleotide sequences into the corresponding amino acid sequences. We downloaded 35,624 human gene nucleotide sequences from CCDS_nucleotide.current.fna.gz and their corresponding amino acid sequences from CCDS_protein.current.faa.gz from the NCBI CCDS database, available via FTP at https://ftp.ncbi.nlm.nih.gov/pub/CCDS/current_human/. We randomly selected 100 sequence pairs to construct the prompt questions and gold-standard answers. To create the question-answering tasks, each prompt included the instruction “Nucleic acid sequence of a human gene:” followed by the nucleotide sequence, and concluded with “Translate the above nucleic acid sequence to the corresponding amino acid sequence.” The corresponding amino acid sequence was used as the gold standard. A score of 1 was assigned if the amino acid sequence generated by the LLM exactly matched the gold standard; otherwise, a score of 0 was assigned.

### DNA sequence extraction

This module evaluates the ability of LLMs to extract the DNA sequence corresponding to a given genomic location on a human chromosome. The DNA sequences and genomic regions used were the same as those in the *DNA sequence alignment to human genome* module. Prompts were constructed by prepending the instruction “Use NCBI database to answer the following question: What is the DNA sequence located at” followed by the genomic coordinate in the format “chr:start-end” (e.g., chr8:7081648-7081782). A score of 1 was assigned if the DNA sequence returned by the LLM matched the gold standard; otherwise, a score of 0 was assigned.

### Develop SeqSnap, a GPT-based application

We developed a GPT-based tool, SeqSnap (https://chatgpt.com/g/g-67c52efdc210819190a9532f264ec9c0-seqsnap), using OpenAI’s GPT Builder platform. This app was created to demonstrate the feasibility of using API-driven execution to tackle sequence alignment questions more effectively. It can recognize when to call external genomic databases (e.g., the National Center for Biotechnology Information (NCBI) database) rather than attempting to generate answers purely from its training data. The concept of integrating the NCBI APIs was described in GeneGPT^17^.

There are two versions of the SeqSnap. The API based SeqSnap requires an OpenAI API key along with access to the NCBI Entrez Programming Utilities (E-utilities)^27^ and BLAST APIs^28^. The workflow begins by using the OpenAI API to classify the user’s question into specific categories. If the task involves aligning sequences, the BLAST API is called with the nucleotide sequence to perform the alignment. For SNP-related queries, the NCBI E-utilities API (specifically the `ESummary`) is used to search the SNP database for relevant information. When a user requests a sequence based on genomic coordinates, the `EFetch` of the NCBI E-utilities API is used to retrieve the sequence. After obtaining the results from BLAST or SNP searches, the OpenAI API is called again to summarize the results into an easy-to-understand format. This system requires packages including `OpenAI`, `pandas`, `requests`, `time`, and `JSON`.

The GPT-based version of SeqSnap follows the same structure, but is designed to be easier to use. Instead of requiring manual API calls, it uses the ChatGPT interface, providing a more intuitive way for users to interact. The integration with the NCBI BLAST and E-utilities APIs allows SeqSnap to perform sequence alignments or search the NCBI database as requested, all through natural conversation. It is easily accessible for users to query genomic data and perform bioinformatics tasks.

### LLMs

The BioGPT and BioMedLM models were accessed using the interface provided by Hugging Face^29^. GPT-3.5 (gpt-3.5-turbo-0125) was accessed via the API provided by OpenAI. Claude 3.5 (claude-3-5-sonnet-20240620) was accessed via the API provided by Anthropic. GPT-4o without online access (gpt-4o-2024-05-13) was accessed via the API provided by OpenAI. Gemini Advanced was accessed through its web browser version (https://gemini.google/advanced/). GPT-4o with online access was accessed through its web browser version (https://chat.openai.com/). GeneGPT (https://github.com/ncbi/GeneGPT)^17^ was implemented using GPT-3.5 Turbo (gpt-3.5-turbo-16k). The slim model used Dm.1 and Dm.4 documents for in-context learning, while the full model used all available documents and demonstrations.

## Data and Code Availability

The full GeneTuring benchmark, including all 1,600 question-answer pairs, prompts to the models, model-generated responses (each question has three replicated queries’ responses), manual scores, evaluation scripts, and documentation, is publicly available at https://github.com/Winnie09/GeneTuring. The repository also includes the documentation of our GPT-based tool, SeqSnap (https://chatgpt.com/g/g-67c52efdc210819190a9532f264ec9c0-seqsnap), which was built using OpenAI’s GPT Builder platform. All materials support open and flexible reuse by the research community.

## Key points

- We developed GeneTuring, a comprehensive and systematically curated question-and-answer benchmark designed to evaluate and enhance the genomic reasoning capabilities of large language models (LLMs).
- Using GeneTuring, we evaluated ten advanced LLM settings, including four versions of ChatGPT, and identified GPT-4o with web access as the top performer across most genomic tasks.
- Although GPT-4o achieved strong performance, it exhibited notable limitations in specific tasks such as sequence alignment. We showed that providing access to external reference genomes, implemented through our GPT-s application SeqSnap, substantially improves accuracy in these contexts.
- These results underscore both the potential and the current limitations of LLMs in genomics, and point to the importance of integrating LLMs with domain-specific tools to achieve more reliable and robust genomic intelligence.

## Acknowledgments

Z.J. was partially supported by the National Institutes of Health (NIH) under Award Number R35GM154865. W.H. was partially supported by the National Institute Of General Medical Sciences (NIGMS) of NIH under Award Number R35GM150887.

## Author contributions

W.H. and Z.J. conceived the study. W.H., X.S., X.L., and Z.J. conducted the analysis. W.H. drafted the manuscript. All authors reviewed, edited, and approved the final version of the manuscript.

## References

1. OpenAI. New embedding models and api updates (2024). URL https://openai.com/index/new-embedding-models-and-api-updates/.

2. OpenAI. Gpt-4 system card (2024). URL https://cdn.openai.com/papers/gpt-4-system-card.pdf.

3. Team, G. et al. Gemini: a family of highly capable multimodal models. arXiv preprint arXiv:2312.11805 (2023).

4. Anthropic. Model card and evaluations for claude models (2023). URL https://www-cdn.anthropic.com/bd2a28d2535bfb0494cc8e2a3bf135d2e7523226/Model-Card-Claude-2.pdf.

5. Hou, W. & Ji, Z. Comparing large language models and human programmers for generating programming code. arXiv preprint arXiv:2403.00894 (2024).

6. Hou, W., Qu, Y. & Ji, Z. Assessing large multimodal models for one-shot learning and interpretability in biomedical image classification. bioRxiv (2024). URL https://www.biorxiv.org/content/early/2024/10/08/2023.12.31.573796. DOI 10.1101/2023.12.31.573796. https://www.biorxiv.org/content/early/2024/10/08/2023.12.31.573796.full.pdf.

7. Hou, W. & Ji, Z. Assessing GPT-4 for cell type annotation in single-cell RNA-seq analysis. Nat. Methods 21, 1462–1465 (2024).

8. Chen, Y. & Zou, J. Simple and effective embedding model for single-cell biology built from chatgpt. Nat. Biomed. Eng. 1–11 (2024).

9. Hendrycks, D. et al. Measuring massive multitask language understanding. arXiv preprint arXiv:2009.03300 (2020).

10. Chen, M. et al. Evaluating large language models trained on code. arXiv preprint arXiv:2107.03374 (2021).

11. Kim, J., Wang, K., Weng, C. & Liu, C. Assessing the utility of large language models for phenotype-driven gene prioritization in the diagnosis of rare genetic disease. The Am. J. Hum. Genet. 111, 2190–2202 (2024).

12. Tang, X. et al. Biocoder: a benchmark for bioinformatics code generation with large language models. Bioinformatics 40, i266–i276 (2024).

13. Sarumi, O. A. & Heider, D. Large language models and their applications in bioinformatics. Comput. Struct. Biotechnol. J. (2024).

14. Luo, R. et al. Biogpt: generative pre-trained transformer for biomedical text generation and mining. Briefings Bioinforma. 23 (2022).

15. Venigalla, A., Frankle, J. & Carbin, M. Biomedlm: a domain-specific large language model for biomedical text. https://www.mosaicml.com/blog/introducing-pubmed-gpt.

16. OpenAI. Gpt-4o system card (2024). URL https://openai.com/index/gpt-4o-system-card/.

17. Jin, Q., Yang, Y., Chen, Q. & Lu, Z. Genegpt: Augmenting large language models with domain tools for improved access to biomedical information. Bioinformatics 40, btae075 (2024).

18. Zhang, Y. et al. Siren’s song in the ai ocean: a survey on hallucination in large language models. arXiv preprint arXiv:2309.01219 (2023).

19. Rawte, V., Sheth, A. & Das, A. A survey of hallucination in large foundation models. arXiv preprint arXiv:2309.05922 (2023).

20. Smith, L. et al. Overview of biocreative ii gene mention recognition. Genome biology 9, 1–19 (2008).

21. Wheeler, D. L. et al. Database resources of the national center for biotechnology information. Nucleic acids research 35, D5–D12 (2007).

22. Frankish, A. et al. Gencode reference annotation for the human and mouse genomes. Nucleic acids research 47, D766–D773 (2019).

23. Amberger, J. S., Bocchini, C. A., Scott, A. F. & Hamosh, A. Omim. org: leveraging knowledge across phenotype–gene relationships. Nucleic acids research 47, D1038–D1043 (2019).

24. Liberzon, A. et al. The molecular signatures database hallmark gene set collection. Cell systems 1, 417–425 (2015).

25. Han, H. et al. Trrust v2: an expanded reference database of human and mouse transcriptional regulatory interactions. Nucleic acids research 46, D380–D386 (2018).

26. Huber, W. et al. Orchestrating high-throughput genomic analysis with bioconductor. Nat. methods 12, 115–121 (2015).

27. Sayers, E. A general introduction to the e-utilities. Entrez Program. Util. Help. [Internet]. Bethesda (MD): Natl. Cent. for Biotechnol. Inf. (US) (2010).

28. Altschul, S. F., Gish, W., Miller, W., Myers, E. W. & Lipman, D. J. Basic local alignment search tool. J. molecular biology 215, 403–410 (1990).

29. Wolf, T. et al. Transformers: State-of-the-art natural language processing. In Proceedings of the 2020 conference on empirical methods in natural language processing: system demonstrations, 38–45 (2020).

