## Supplementary Figure for "Benchmarking large language models for genomic knowledge with GeneTuring"

### Supplementary materials

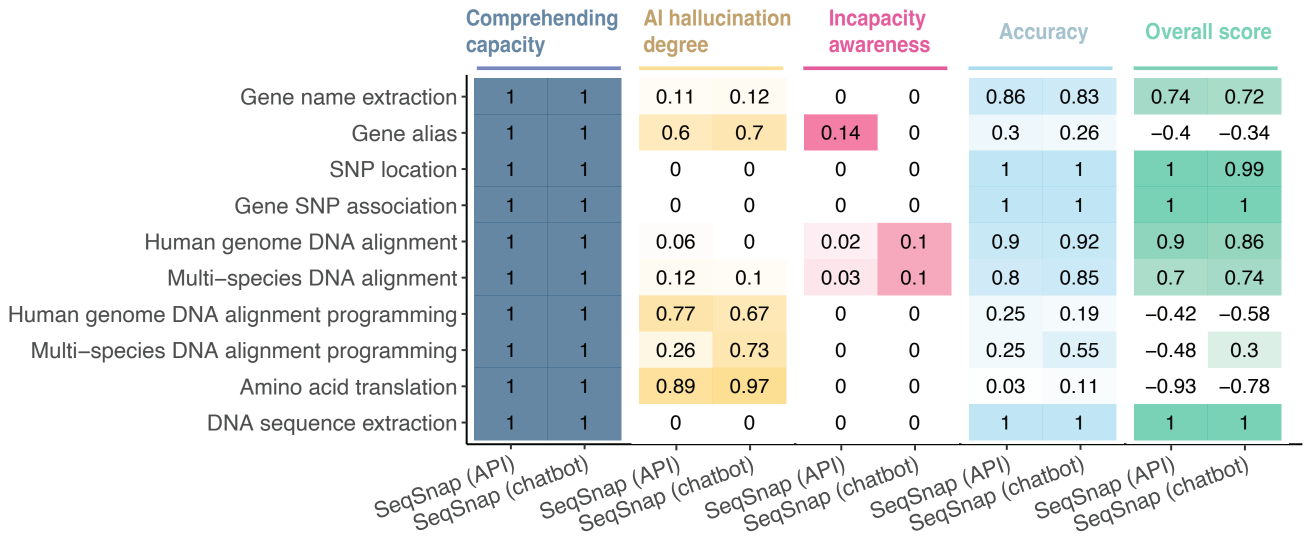


**Figure S1. Performance comparisons of the API and chatbot versions of SeqSnap on GeneTuring modules.** [1]

[1] All questions were assessed for the API version, and the top 10 questions were assessed for the chatbot version due to time constraints.
